# The N-glycosylation sites and Glycan-binding ability of S-protein in SARS-CoV-2 Coronavirus

**DOI:** 10.1101/2020.12.01.406025

**Authors:** Wentian Chen, Ziye Hui, Xiameng Ren, Yijie Luo, Jian Shu, Hanjie Yu, Zheng Li

## Abstract

The emerging acute respiratory disease, COVID-19, caused by SARS-CoV-2 Coronavirus (SARS2 CoV) has spread fastly all over the word. As a member of RNA viruses, the glycosylation of envelope glycoprotein plays the crucial role in protein folding, evasing host immune system, invading host cell membrane, even affecting host preference. Therefore, detail glyco-related researches have been adopted in the Spike protein (S-protein) of SARS2 CoV from the bioinformatic perspective. Phylogenic analysis of S-protein sequences revealed the evolutionary relationship of N-glycosylation sites in different CoVs. Structural comparation of S-proteins indicated their similarity and distributions of N-glycosylation sites. Further potential sialic acid or galactose affinity domains have been described in the S-protein by docking analysis. Molecular dynamic simulation for the glycosylated complexus of S-protein-ACE2 implied that the complicate viral binding of receptor-binding domain may be influenced by peripheric N-glycans from own and adjacent monoers. These works will contribute to investigate the N-glycosylation in S-protein and explain the highly contagious of COVID-19.

## Intrudction

Recent COVID-19 (Coronavirus Disease 2019) caused by a novel coronavirus named Severe Acute Respiratory Syndrome Coronavirus 2 (SARS-Cov-2) has been spread fastly all over the world. Higher lethality and powerful human-to-human transmission capacity has aroused widely concern. As a kind of enveloped virus with singlestranded positive-sensed RNA, new SARS2 CoV (Coronavirus) is a member of CoV family, which has closer relationship to previous SARS (severe acute respiratory syndrome) and MERS (Middle East respiratory syndrome) CoVs ^[1,2]^.

Coronaviruses are cataloged into the Nidovirales, Cornidovirineae, Orthocoronavirinae and divided into four Genuses. Until now, the Alpha-coronavirus and Beta-coronavirus response to known human-isolated CoVs (HCoVS), including the above three CoVs combined with HKU1, OC43, NL63 and 229E HCoVs. According to phylogenetic analysis, these HCoVs are considered to have originated from the bats and rodents ^[3–5]^. The genomes of coronaviruses have been described meticulously in previous reports. As one of biggest viruses, the 5’-terminal of positive-sense and single-stranded RNA (+ssRNA) genome in coronavirus encodes a polyprotein complexus, pp1ab, while the 3’-terminal encodes the structural proteins, such as the envelope glycoprotein spike protein (S-protein), envelope (E), membrane (M), nucleocapsid (N) and possible hemagglutinin-esterase (HE)^[6–7]^.

The clove homotrimeric S-protein is a type I glycoprotein which gives the crown-like appearance on CoVs. The S1 and S2 subunits in S-protein monomer, are responsible for cell binding and membrane fusion, respectively ^[8]^. The S1 subunit forms the globular head and contains the N-terminal domain (NTD), receptor binding domain (RBD) and smaller subdomains (SD1 and SD2). The S2 subunit is conserved among all coronaviruses and forms the main rosette-like α-helix bundle and a β-sheets-riched subdomain. In the endosome, the S2 could be further cleaved by the host proteases and exposed its fusion peptide, which resulting in final membrane fusion ^[9–10]^.

The receptor of ACE2 (angiotensin-converting enzyme 2) for SARS CoV and DPP4 (dipeptidyl peptidase 4) for MERS CoV have been reported ^[11–12]^. However, the S-protein is highly glycosylated, as many as 22 potential N-glycosylation sites in the S-protein of SARS CoV could be detected, compared to 23 N-glycosylation sites in the S-protein of MERS CoV ^[13–14]^. Therefore, it is worth to figure out the distribution of N-glycosylation sites and glycobiology functions in the S-protein of SARS2 CoV.

In this article, we have compared the evolutionary relationship and distribution of N-glycosylation sites in different CoVs. For the distribution and possible functions in the N-glycosylation sites, a homologous modeling method had been adopted. Further docking analysis have provided a visual method for the possible glycan binding domains. These works should contribute to explan the highly contagious of new SARS2 CoV and provide new strategy on SARS2 CoV prevention.

## 1 Methods

### 1.1 Phylogenetic analysis for S protein and N-glycosylation sites

For the purpose of the evolution of S-protiens and N-glycosylation sites in CoVs, a dataset of S-protein sequences from representative 1169 CoVs was retrieved from the National Center for Biotechnology Information (NCBI) VIRUS database (https://www.ncbi.nlm.nih.gov/labs/virus, accessed in April.15th 2020), containing approximately 438 human reports in 1023 sequences^[15]^. An alignment of whole sequences was performed by ClustalW 2.0 in MEGA 7.0 (File S1)^[16]^. A webtool named “NetNGlyc 1.0 Server” was used for the N-glycosylation sites predicting^[17]^. To investigate the evolutionary relationship of N-glycosylation sites in different subgenus, a smaller dataset was used for further analysis with 49 representative sequences, including the HKU1, OC43, NL63, 229E, SARS, MERS as well as SARS2 CoVs. Unrooted phylogenetic tree was constructed using the Neighbor-Joining method and the Poisson correction model. The internal branching probabilities were determined by bootstrap analysis with 1,000 replicates similar to previous description^[18]^.

### 1.2 Homologous modeling for S-protein of SARS2 CoV

As the SARS2 CoV continued to spread across the world, more and more viral complete genomes were sequenced, as well as the crystal structures of S-proteins. However, the available structures are lock of partial peripheral elements^[19]^. Hence, the amino acid sequence from one S-protein (Genebank ID: QHN73810.1) was selected for homologous modeling by SWISS-MODEL web service^[20]^. As a result, a 3D structure file of S-protein with the largest similarity of 99.9 % to a reported S-protein of SARS2 CoV (PDB ID: 6VSB) was created^[21]^. For the structural optimization, a 10 ns molecular dynamic (MD) simulation has been adopted. Minimization and equilibration were performed using the NAMD 2.8 program with the CHARMM22 all-atom force field for the protein as previous describing^[22]^.

### 1.3 Molecular docking analysis for Glycan ligands candidates and S-proteins

In order to assess the possible glycan-binding ability of S-protein of SARS2 CoV from the virtual calculated perspective, a serial of glyco-ligands were designed, including the monosaccharides (The Man (Mannose), Gal (Galactose), Glu (Glucose),GalNAc (N-acetyl-β-D-Galactosamine), GlcNAc (N-Acetyl-β-D-Glucosamine), Xly (Xylose), Fuc (Fucose) and SA (Sialic Acid)), and common terminal structures at the N-glycans, such as the disaccharides (G3GN(Galβ-1,3GlcNAc),G4GN (Galβ-1,4GlcNAc), GN2M (GlcNAcβ-1,2Man), GN4M (GlcNAcβ-1,4Man), SA23Gal (SAα-2,3Gal), SA26Gal (SAα-2,6Gal)). The Blood group antigens (Blood group A(Fucα-1,2Gal[β-1,4GlcNAc]β-1,3GalNAc), Blood group B antigens (Fucα-1,2Gal[β-1,4GlcNAc]β-1,3Gal), Blood group o antigens (Fucα-1,2Galβ-1,4GlcNAc) and other comparative saccharides such as the X2X (Xylβ-1,2Xyl), G4G (Glcβ-1,4Glc), X4X4X4X (Xylβ-1,4Xylβ-1,4Xylβ-1,4Xyl), MMM (Manα-3,6Manα-3,6Man), 3-sialyllactose (SAα-2,3Galβ-1,4Glc), 6-sialyllactose (SAα-2,6Galβ-1,4Glc), Disialyllacto-N-tetraose (DSLNT). All of the glyco-ligands were constructed by the online SWEET-H program and optimized by the MM3 force field^[23]^.

The S-protein model of SARS2 after MD simulation was subjected to docking analysis. The automated docking analysis between S-protein and glyco-ligands was performed using the AutoDock Vina program. A grid box with size of 70 × 70 × 70 points grid box was used to cover the mainly top surface of S-protein monomer during the docking analysis. The receptor atom positions were held fixed, and the glycosidic bonds of saccharides were variable. Other docking parameters were set to default ^[24]^. As the contrast, the S-proteins from SARS (PDB ID:6ACC), MERS (PDB ID: 5W9J), NL63(PDB ID: 5SZS), MHV CoVs (PDB ID: 3JCL), were selected for the same procedures.

### 1.4 The N-glycan analysis in the S-protein-ACE2 interaction

To figure out whether N-glycans from both S-protein and ACE2 impact the receptor-ligands interaction, the further MD simulation for glycosylated ACE2-S-protein complex were performed based on the reported coordinated file (ACE2-NTD: 6M0J; Up-stand stated S-protein: 6VSB). The ACE2-NTD PDB file, which contains the ACE2 and incomplete S1 subunit was subsequently superposed by the optimized SARS2 model and up-started S-protein respectively. The “complex” type N-glycans, were added on all N-glycosylated sites by “glycoprotein builder” program in GLYCAM online server^[25]^. An NVT ensemble (canonicalensemble) and a revised force-field file for glycoprotein (File S2) was used in the MD simulation for these complexus, further distance analysis for the fluctuations of N-glycans was adopts by VMD1.9.3^[22]^. Five pairs between the geometric center of one GCD1 to the terminal SA and Gal residues, the inner Man, GlcNAc residues and glutamic acid residues from N90 in ACE2 were selected for possible interaction assesssing. The distance of “GCD1-N90, GCD1-GlcNAc, GCD1-MAN, GCD1-Gal and GCD1-SA” were sampled every 100 ps during 10 ns MD simulation.

## 2 Results

### 2.1 The pholygenic analysis of the S-proteins in CoVs

There are four genuses in Orthocoronavirinae, included the Alpha-, Beta-, Delta- and Gamma-coronaviruses. The Betacoronavirus genus could be further divided into the Embecovirus, Hibecovirus, Merbecovirus, Nobecovirus, Sarbecovirus and others subgenuses^[26][25]^. Unlike the continuous-spreading the IVs (Influenza Viruses) and HIVs (Human Immunodeficiency Viruses), most of records were derived from outbreak epidemic CoVs, while their amino acid sequences are relative conservative.

Based on the genome similarity, the emerging SARS2 CoV showed the minimal difference to the BAT SARS-like CoV (e.g. Genebank ID:AVP78042.1, within 99% sequences similarity) and clustered in the Sarbevirus subgenus^[27]^. Up to 25^th^ June 2020, more than 90 SARS2 CoV genomes have been included in the NCBI database. The similarity of the S-protein amino acid reaches up to 99.9% and few residues are mutated in 1273 amino acid full-length (e.g. H49Y and S247R mutations. The numbering system of S-protein of SARS2 CoV was in keeping with an easier reported sequence. Genebank ID: QHN73810.1)^[28]^).

The genome of CoVs varies from 28 k to 31 k bp (base-pair), and the length of S-protein differs from 1100 to 1500 aa (amino acid. e.g. 1457 aa in Canine coronavirus, Genebank ID: BAW32706.1). Up to 1th June, nine CoVs have the more than 100 records. The SARS2 (4130 records), SARS (303), MERS (734), OC43 (291), BCoV (245) in Beta-coronavirus, IBV (Infectious bronchitis virus, 580) in Alpha-coronavirus, PDC (Porcine Deltacoronavirusvirus, 223) in Gamma-coronavirus, the PEDV (Porcine epidemic diarrhea virus, 1890) and 229E CoVs (101) in Delta-coronavirus as well as other representative CoVs are selected for further analysis.

The N-J tree of S-protein derived from the MEGA7 is similar to those from the whole-genome researches (Figure 1A)^[29]^. But the sequence similarity of various S-proteins, even within the Betacoronaviruses, is rather low. The homology of S-protein from Sarbecovirus subgenus, which included the Bat SARS-like, SARS and SARS2 CoVs in a 338 analytical dataset, reaches up to 57.8%, and most conserved residues distribute at the C-terminal (File S3).

**Figure1.**
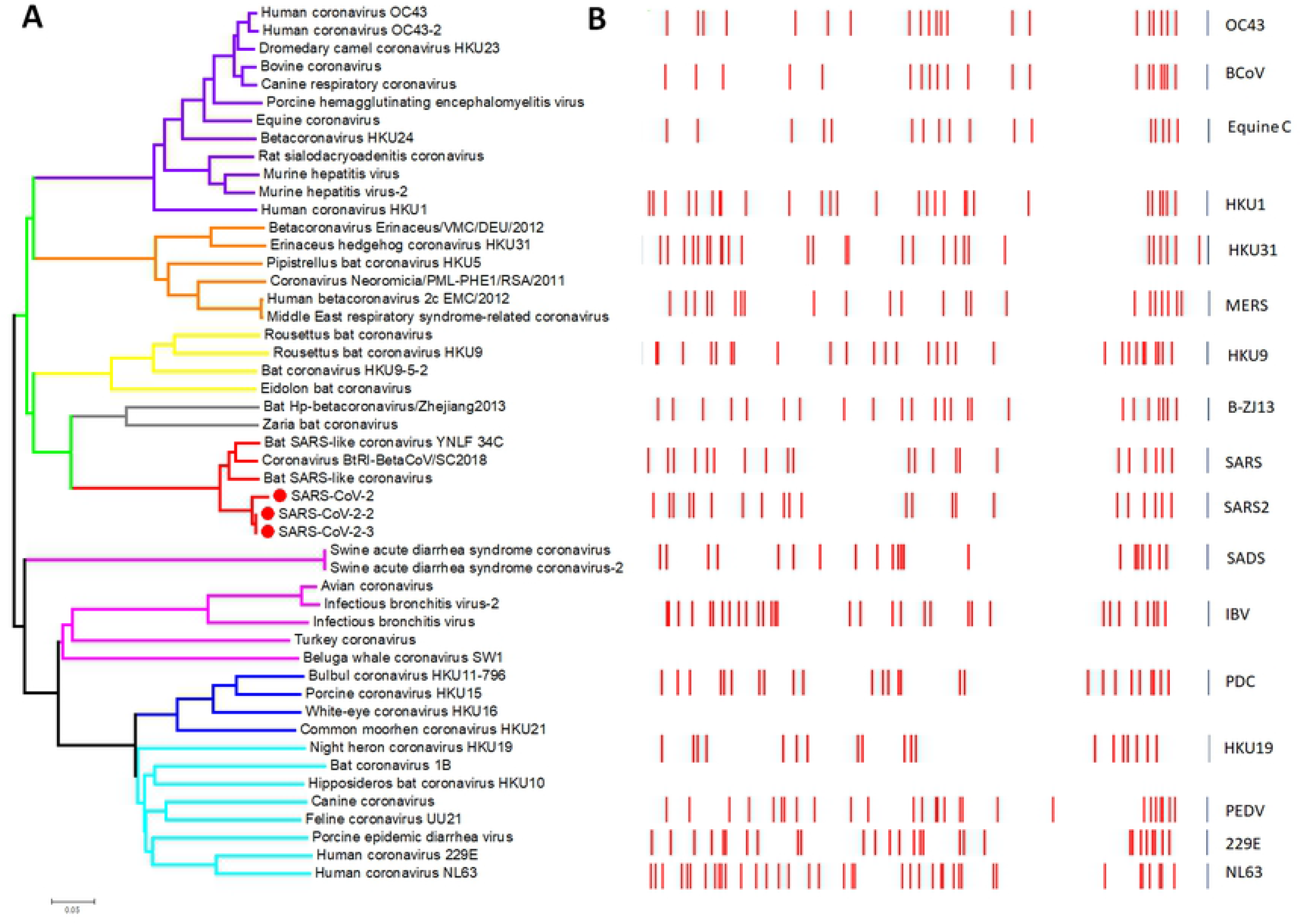
The N-J Phylogenetic tree and the distribution of N-glycosylation sites in S-proteins from the representative CoVs. (A). More than 50 S-protein sequences from different CoVs are selected for the phylogenic analysis. Current S-proteins can be classified into different clades. The purple, green, blue and cyan clades denote the Alpha-, Beta-, Gamma- and Delta-coronaviruses respectively. The indigo, orange, yellow, dark grey and red subclades in the green clade represent the Embecovirus, Merbecovirus, Nobecovirus, Hibecovirus, Sarbecovirus subgenus. The S-proteins of emerging SARS2 CoV are labeled as the red spheres. (B). The distribution of N-glycosylation sites from different S-proteins are shown in bar charts. Although the lengths of S-proteins varied greatly, all the slides are set to the same length for observing the distribution of glycosylation sites. The abbreviations are consistent with the level CoVs.

### 2.2 Comparing the structures of S-protein

Up to now, the reported S-protein structures of SARS2 CoV are partial missing, such as the peripheral elements at the NTD (e.g. PDB ID:6X29, 6VSB). The “SWISS-MODEL” provided one proper method for the S-protein reconstruction. Based on the existing structures, the created model has an identity of 99.7% to a reported S-protein from SARS2 CoV^[30]^. After a 10 ns MD simulation for S-protein, the optimal model was used for structural comparation to those from PEDV (PDB ID: 6U7K) in Alpha-coronaviruses, SARS, MERS, and MHV (Mouse hepatitis virus, PDB ID: 6VSJ) in Beta-coronaviruses, HKU15 (Porcine coronavirus HKU15, 6B7N) in Deltacoronavirus and IBV (PDB ID: 6CV0) in Gamma-coronavirus, all S-proteins share the similar structural characteristics but with larger differences in the S1 subunit, which is in accord with above sequence analysis (Figure S1).

Three S-protein monomers twist together and form the obconic (or clove shape) trimmer, while each monomer consists of global S1 and stalk S2 subunit (Figure 2A). Similar to the description from other coronaviruses, in SARS2 CoV, the S1 subunit can be divided into the NTD (N-terminal domain,1-290), RBD (Receptor-Binding Domain, 330-528), SD1 (subdomain1, 320-329 and 530-590.) and SD2 (subdomain2, 309-322 and 589-653) in order^[31]^. The stalk S2 subunit participates in the virus-host membrane fusion and consists of conserved bunches of α-helixes (728-1069). In addition, a third subdomain named SD3, as well as the TM (Transmembrane) and CP (Cytoplasmic tail), at the C-terminal (1070-1273), are partial avaible in all identified structures.

**Figure 2.**
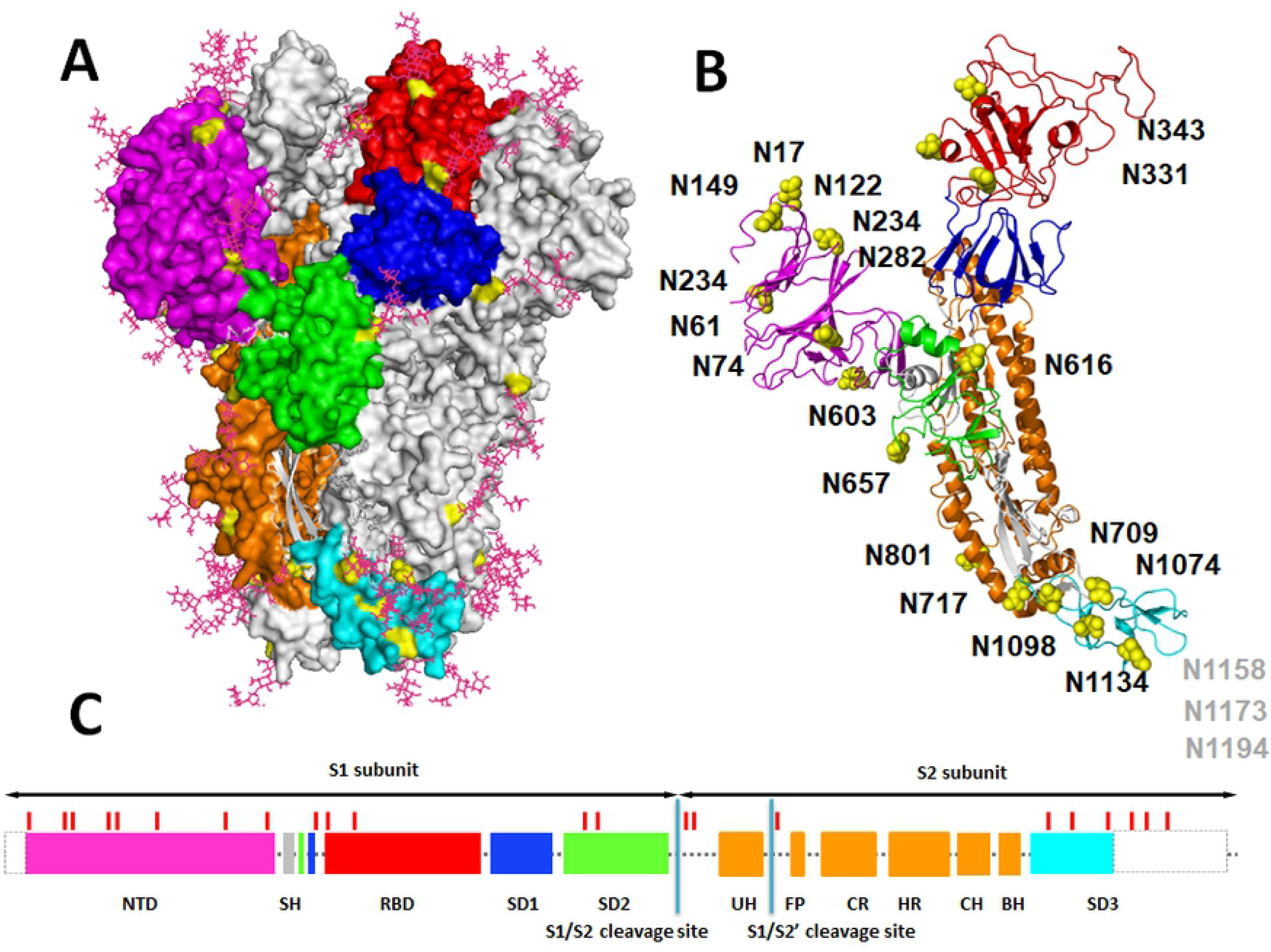
The structure of S-protein of SARS2 CoV. (A). The S-protein is a clove trimer and highly N-glycosylated. The S-protein monomer can be divided into NTD (purple), RBD (red), SD1 (blue), SD2 (green), α-Helix-bundles (orange), SD3 (cyan) and so on by structural characters. The N-glycosylation sites are denoted in yellow, while the N-glycans are show in the purple sticks. (B). The distribution of N-glycosylation sites in the monomer. The C-terminal N1158, N1173 and N1194 are not shown in the missing element. (C). The diagram of S-protein describes the sequential subdomains and the location of N-glycosylations. Undected regions in the coordination file are shown in dash blocks.

As is shown in Figure 2B, NTD from SARS2 CoV are located at the triangular angles. A typical β-sandwich core structure co-exists in all S-proteins, which is consisted of one 5-stranded β-sheet and one 6-stranded β-sheet in SARS2 CoV^[32]^. A smaller helix element (291-308) co-exists in all S-proteins and responsible for NTD and RBD connecting. The RBD has been described detailedly in different CoVs, such as the flexible RBD of SARS and SARS2 CoVs are readily recognized by its receptor with either lying and Up-standing state^[33,34]^. The precise coordinate files of RBD-ACE2 interaction compleuxs from both SARS and SARS2 CoVs had been reported. We found that a Tyr-riched region at the interactive interface of RBD was also relative conserved in other Beta-coronaviruses (Figure S2). Noteworthy, only one RBD in S-Protein of SARS and SARS2 participate in the ACE2 binding, and the mechanism needs further study^[35]^. Two smaller β-sheets-riched SD1 and SD2 appear to be the base to underpin the NTD and RBD. Interestingly, these subdomains are composed of two discrete sequences (Figure 2C). The S1 and S2 subunits are linked by the S1/S2 cleavage site, and “RRAR” motif in SARS2 CoV is regarded as the furin recognition site, rather than fewer basic residues in other coronaviruses^[36]^. The rosette-like S2 subunits are consisted of bunches of α-helixes which can be further devided into smaller helix elements. Moreover, it can be inferred that uncompleted SD3 is rich of β-sheets.

### 2.3 N-glycosylation sites in S-proteins of SARS2 CoV

S-proteins are highly N-glycosylated, commonly, as many as 20 potential N-glycosites can be detected from different CoVs. There are 22 potential N-linked glycosites in the S-protein of SARS2 CoV, while SARS-CoV and Bat SARS CoV possess 22 and 23 glycosites respectively (Figure1B). In all CoVs, the numbers and distribution of sites is not closely related to the genuses. As is shown in Figure 1B, the N-glycosylated sites of representative CoVs mainly cluster in the S1 subunit and the C-terminal of S2 subunit. Although the length of S-protein is different, the distribution of N-glycosylate sites is similar. However, unlike the continual IVs and HIVs, which contain abundant emerging or missing N-glycosylation sites, most glycosites in CoVs are conserved^[18]^.

It is well-known that, the N-X-S/T (X can not be Pro residue) sequon is the motif for N-glycosylation. In SARS2 CoV, eight potential N-glycosites distribute in NTD (including N17, N61, N74, N122, N149, N165, N234 and N282). It reflects that the NTD is under continual host immune surveillance, while N-glycans may shield the epitopes. Only one N-glycosite, N343, locates at the top of RBD and near to the geometric center of the trimerical top. Another N-glycosite, N331 is at the linker of RBD and SD1, followed by N603, N616 and N657 in SD2. A long linker region connects the S2 cleavage site to the long upstream helix (UH). N709 is near to the S1/S2 protease cleavage site, and highly conserved in ases^[37]^. Another similar N-glycosite, N801, may participate in the protection of the S1/S2’ protease cleavage site. The rest of N-glycosites, including N717, N1074, N1098, N1134 and invisible N1158, N1173, N1194 in SD3 locate at the bottom of S-proteins (Figure 2C).

### 2.4 The glycan-recognition domains in the S-protein of SARS2 CoV

The crystal structures of RBD-ACE2 complexus had been reported from the lastest works^[38,39]^. We found that eight-Tyr residues in the RBD form the unbond-interactions with the N-terminal helix fo ACE2**Error! Reference source not found.**. By comparing this interactive region to other CoVs, most of Tyr residues are conserved in Serbevirus subgenus (Figure S2)^[40]^. Although protein-protein interactions are also reported in other CoVs, such as the DDP4-MERS or CEACAM1a-MHV^[41]^, in view of present works, glycan-protein recognization is also universal in viral invasion, such as the sialic acids (SA) residue bind to Hemagglutinin in Type A IVs^[42]^, the N-acetyl-9-O-acetylneuraminic acid binds to HE (Hemagglutinin-Esterase) in Type C IVs, the S-protein of HKU1 or BCoV^[43]^. Hence, an explorative work on the potential glycan-recognition domains (GCDs) in S-protein of SARS2 has been took by using docking analysis.

Five different S-proteins had docked with a series of glyco-ligands, which included common sialoglycan ligands (SA, SA23Gal, SA26Gal, 3-sialyllactose and 6-sialyllactose.), gal-related lignds (Gal, SA23Gal, SA26Gal, G3GN and G4GN) and others (monosaccharides such as the Glc, Xyl or the X4X4X4X and MMM.). As listed in the Figure 3, the binding energies derived from Autodock Vina results indicate different binding abilities of these S-proteins by theoretical calculation. The lowest binding energy (best binding ability) comes from the NL63-3-sialyllactose (−9.4 Kcal/mol) while the highest from the SARS2-X4X4X4X and NL63-Fuc (−4.2 kcal/mol). Taken together, sialoglycan and Gal related-glycans showed higher binding energies to SARS2 model. In view of different blood type, the S-protein of SARS2 did not show obvious preference to these Gal- and GalNAc-terminal blood group antigens. These results may reveal the S-proteins binding preferences.

**Figure 3.**
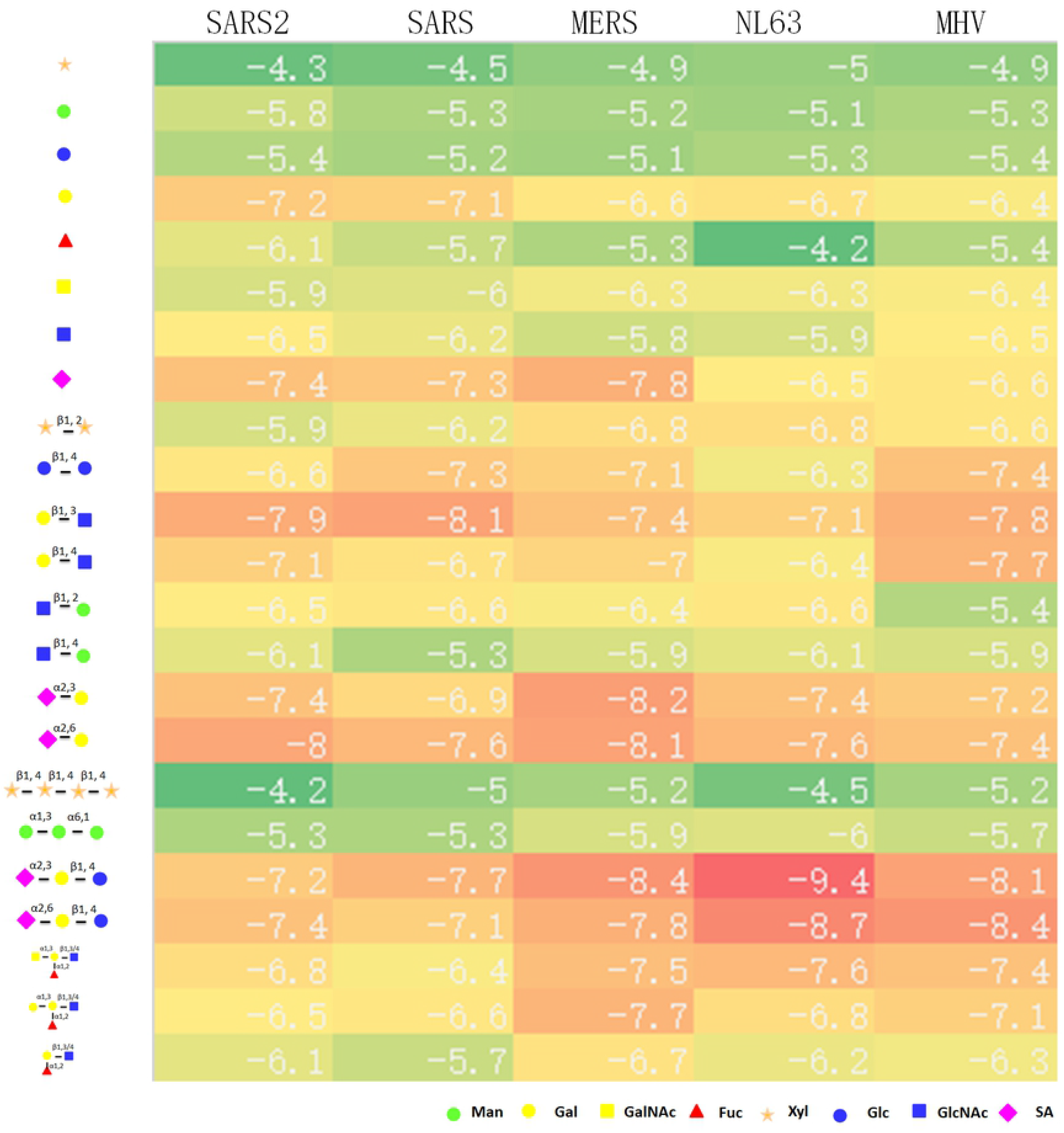
The predicted docking energies of S-proteins and saccharide ligands. As is listed in the left line, all test ligands are shown in the symbolic representation mode. The color lumps in the heatmap indicated the magnitude of the binding energies. Interestingly, the SA- or Gal-terminal saccharides show higher binding abilities to S-protein of SARS2 CoV.

The ADT provides the visual methods for observing the receptor-ligand interactions. In these docking assays, three possible glycan-recognized domains (GCDs) with higher binding ability could be concluded in the S-protein of SARS2 CoV. These potential GCDs are close to the NTD and surrounded by peripheral small helix, β-sheet or Loop. As is shown in Figure 4, first potential GCD (GCD1, red region), which appears at the top of NTD, has showed stronger affinity to SA-related and Gal-related glycans. It locates at the top of β-sandwich core and constitues of one smaller helix, two samller β-sheets and long loops. The crucial residues, E155, V157, K115 and V113 on the bottom β-sheets and the Y146, W138 at the Helix140 participate the H-bonds formation in the binding pocket. The SA residues in sialoglycans, as well as the Gal residues in Gal-related saccharides, adopt lying conformation and form the H-bonds with peripheric residues. Similar to the NTDs from other CoVs, a typital β-sandwich core structure in SARS2 CoV consisting of one 5-stranded β-sheet and one 6-stranded β-sheet (Figure S3)^[32]^. Previous studies hinted NTD in MHV or OC43 CoV have affinity to N-acetyl-9-O-acetylneuraminic acid, and their NTD showed high similarity to human galactose-binding lectin domain (e.g. Galectin-4. PDB ID: 5DUU)^[44]^, which also contain peripheral structural elements, mostly long loops and short-sheets, on top of the core structure (Figure 4C).

**Figure 4.**
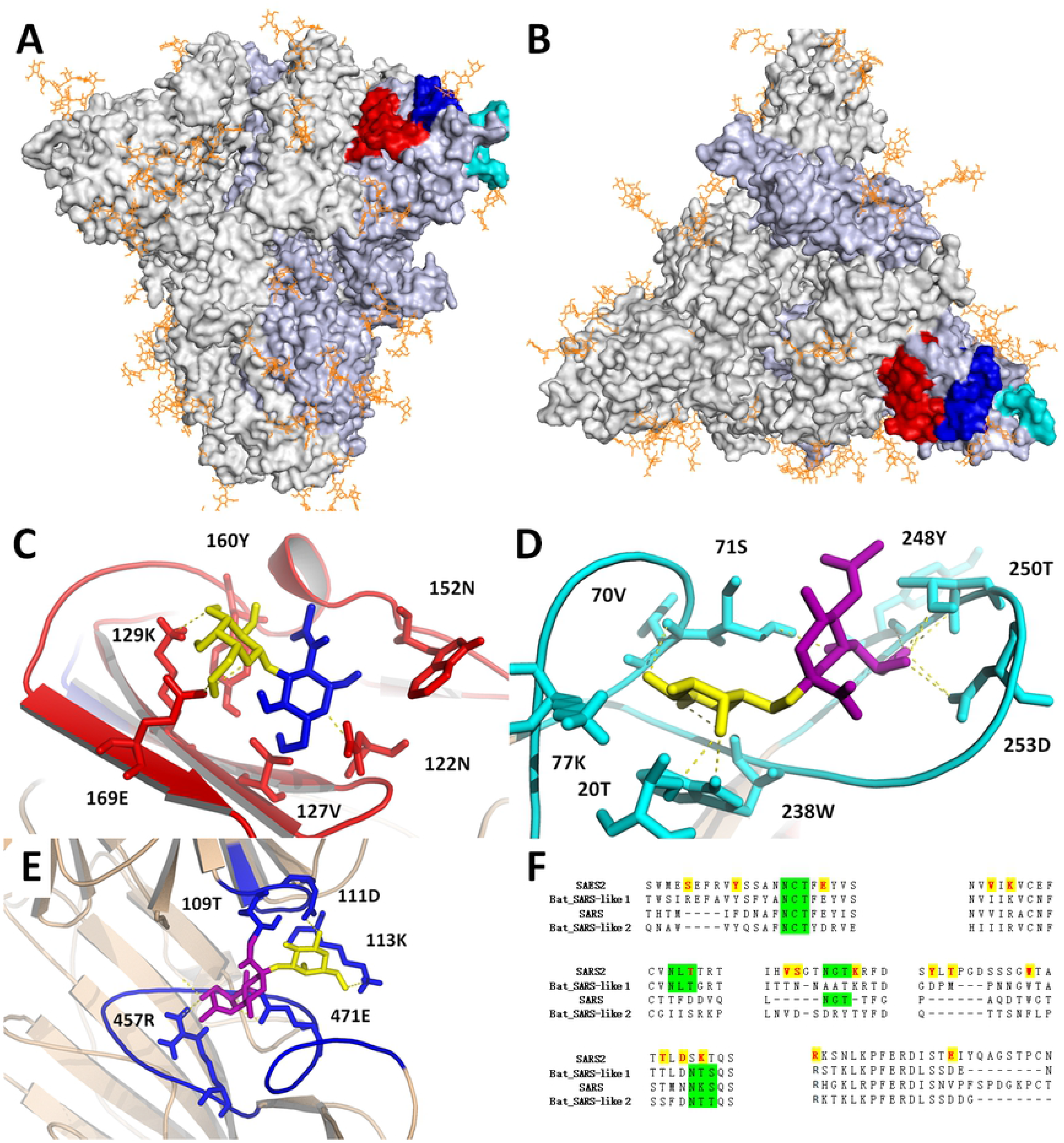
The potential glycan-recognition domains (GCDs) in the S-protein of SARS2 CoV. (A, B, C). Three possible GCDs in S-protein distribute in the NTD. The GCD1-3 are shown in the colored cartoon modes while the key residues revealed in the stick models. The blue, yellow, and purple residues in the docking center represent the GlcNAc, Gal and SA residues respectively. (D). The alignment of GCDs to the counterparts from other members in serbevirus subgenus. SARS2 (NCBI NP: YP 009724390.1), Bat SARS-like 1(AAZ41329.1), SARS (ABD72993.1), Bat SARS-like 2(AAZ41329.1). The Key residues in the ligand-receptor are shown in the red color while the N-glycosylation sites are labeled in the green.

The GCD2 locates at the trimerical outermost angles (Cyan) and shows higher binding affinity to most sialoglycans, especially to the SA23GAL. This domain is consisted of the Loop10, Loop250 and Helix150, while the V16, S254 and E154 may play the key roles in receptor-ligand interaction (Figure 4D). Interestingly, this domain have also been discussed in other CoVs such as the MHV, OC43 and NL63 CoVs^[45–47]^.

Parts of sialoglycans, like SA26GAL in MERS, also show higher affinity to the GCD3 (Blue). This domain locates between the GCD1 and GCD2. The key residues in GCD3, like T109, D111, K113 and R457, are distinctive in SARS2 CoV, where one miss N-glycosylation sites (N254S) also can be found in either SARS or Bat-like SARS CoVs (Figure 4 E,F).

However, all docking tests were performed without consideration of N-glycosylation. It is worth nothing that one or more N-glycosylation sites located at the edge the above GCDs. Such as the N165 in GCD1, N18 and N74 in GCD2.

### 2.5 N-glycan Effects of S-protein and ACE2 interaction

Previous studies in SARS CoV had elaborated the binding mechanism between the S-proteins and ACE2. Only one “up” RBD in trimeric S-protein binds ACE2 molecular by using a protruding up conformation^[48,49]^. More than eight Tyr residues in RBD participated in the nonbonded interaction with the N-terminal helix of ACE2. This Tyr-riched region locates at the top of trimmer and faces to the outside, which is consisted of two bands β-sheets bottom and surround loops. Given the structural similarity of S-protein between SARS and SARS2 CoVs, it seems likely that the similar binding mechanism also occurs in the SARS2 CoV, while the emrging F489Y mutation also increase its binding affinity (Figure S2). Acturally, the latest researches from the complexus of ACE2 and RBD in SARS2 CoV verified this hypothesis ^[30,33,34]^.

However, the N-glycosylation in this complicated interaction is usually neglected. ACE2 is also highly glycosylated, including six N-glycosylation sites and possible O-glycosylated region^[50]^. The incomplete ACE2-RBD complexus in PDB database also denotes that five N-glycosylation sites (N53, N90, N103, N322, N546) in ACE2 distribute around the interactive interface (PDB ID: 7BZ5, 6M17, 6M0J, 6LZG, 7BWJ, 7C01, 6VW1 and so on.)^[51,52]^. Acturally, as many as fourteen N-glycosylation sites, including five in ACE2, two in RBD (N343, N331) and eight in nearby monomer (N122, N165, 2*N234, N331, 2* N343), distribute surround the interactive interface (< 50 Å, Figure S4). Among of these, N322, N90, N122 in ACE2, and N165 in RBD with the distance to the RBD center shorter than 30 Å.

As observed from the existing ACE2-RBD complexs derived from SARS or SARS2 CoVs, we found that the N-terminal helix of ACE2 laying on the RBD and orienting to the adjacent NTD. This would result the N-glycan at the N90 of ACE2 appear on the adjacent NTD and the terminal residues of N-glycan interact with the GCD1 directly (Figure 5). By considering the SA- and Gal-affinity of GCD1, whether an unknown recognition mechanism may promote the ACE2-S-protein binding aroused our interest.

**Figure 5.**
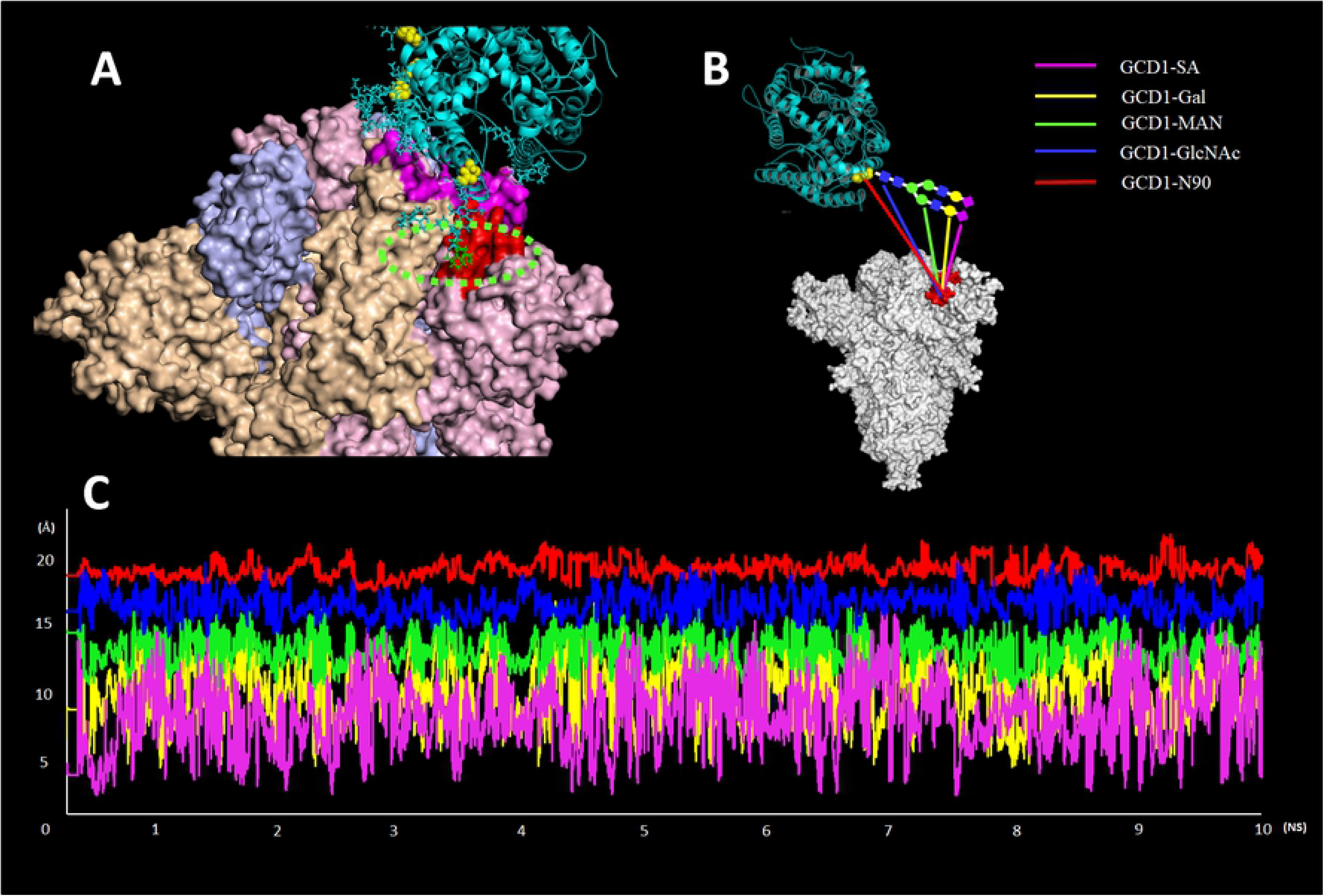
The N-glycan of N90 in ACE2 may interact with the GCD1 of S-proteins. (A). At the foremost step of viral invasion, the N-terminal helix of ACE2 and RBD form the compact interaction, meanwhile, the terminal residues in N-glycan (green dashed oval) of N90 of ACE2 may contact the GCD1; (B). During a 10 ns MD simulation, the centers of terminal residues in the N-glycan of N90 in ACE2 to GCD1 fluctuate between 2 to 15 Å. It hinted that another possible glycan-protein binding mechanism may exist in the SARS2 invasion.

In order to discuss the influence of N-glycans in ACE2-RBD interaction, glycosylated ACE2-S-protein complex was built by the GLYPROT online serve. Based on N-Glycosylation researches, the “complex type” N-glycans were added to the N-glycosites in ACE2 and S-protein. The distance analysis from the “GCD1-N90, GCD1-GlcNAc, GCD1-MAN, GCD1-Gal and GCD1-SA” pairs also indicated the close contact between the N-glycan and one GCD1. During the 10 ns MD simulation, the N-glycans swing around their glycosites and the terminal residues in N-glycans fluncated more flexible. By overlapping the RBD of RBD-ACE2 complex to the same domain of S-protein trimer by either laying or standing conformation, Obviously, the N-glycans from the N90 of ACE2 may contact the GCD1 frequently. It hints that during the viral invasion, the dual binding mechanism may exist in the ACE2-S-Protein interaction, multi Tyr residues may form stronger nonbonded interaction while weaker binding affinity from the Gal- or SA-terminal of N-glycan to NTD may enhance this binding ability.

## 3 Discussion

### 3.1 The distribution of N-glycosylation sites in coronaviruses

Glycosylation plays an important role in the viral life cycle, and N-glycosylation is necessary for viral envelope glycoproteins, such as the nascent glycoprotein folding, maturation, or degradation, escaping host immune surveillance, regulating the sensitivity to temperature adaption, protection of cleavage sites, even impacting pathogen-host interaction^[53]^. As the most striking protein in CoVs, S-protein is one of glycoproteins (M-protein is another glycosylated protein, which possess one to six conserved N-glycosylation sites in different CoVs and consisted of 1100-1400 amino acids, File S4.)

Seldom glycoprotein contains so many N-glycosylation sites like S-protein, while 39 glycositescan be found in the S-protein of NL63 CoV (NCBI GENEID: AWK59943.1). It is can be concluded that more than one hundred of N-glycans in this trimer S-protein and the mass glycans surround membrane matrix. Previous works on IVs have found that most of N-glycosylation sites in the N-terminal of HA (or HA1) varied largely in different subtypes, while the glycosites in stem HA2 domain are highly conserved, this is similar to S-protein. Analogously, the sequence similarity of CoVs in different genues is lower, but their global structures are conserved. As many as 20 glycosites can be detected in different S-protein monomer, while most of N-glycosites scatter at the S1 subunit and 6 to 8 conserved N-glycosites cluster at the C-terminal. Large differences of glycosites exist among four genuses, even inside the Betacoronavirus genus. By comparing the glycosites in Sarbecovirus subgenus, both 22 glycosites cound be found in SARS and SARS2 CoV, while 23 glycosites in BAT SARS-like CoV (File S4).

Although there are numerous N-glycosylation sites in S-protein, but whether they are all glycosylated? Kelley WM *et al*. pointed that the nascent 14-sugar glycan transferred to the N-X-T/S sequon in endoplasmic reticulum firstly, subsequent glycan processing is along with the peptide folding^[54]^. Actually, the S-proteins included in the PDB database also verified the highly glycosylated. The GlcNAc residue at the Asn spread all over the surface of S-protein (e.g. MERS (PDB ID: 5W9P), SARS (PDB ID: 5X58) or SARS2 CoVs (PDB ID: 5W9P)), which is the feature of the N-glycan cleavaged by the PNGase^[55,56]^. What’s more, LC/MS provided the exact N-glycan structures at 9 glycosites from NTD of IBV in Parsons’ works, including the “high-mannose”, “complex” and “hybrid” glycans^[57]^. The deletion of 6 glycosites results the losing of binding ability. These results are consistent with the N-glycosylation in SARS CoV^[58]^. According to the latest analysis, both N-glycans isolated from SARS and SARS2 CoVs are similar, namely including “high mannose”, “hybrid”, “complex” N-glycans. Interestingly, the oligomannose-type glycans were abundant at the bottom of S-protein and the N234 which near to the RBD binding domain.Across the 22 N-linked glycosylation sites, 16% of the glycans contain at least one SA residue and 48% are fucosylated^[59]^.

In this study, S-protein is roughly divided into NTD, RBD, SD1, SD2, Helix-Bundles and SD3 from the N- to C-terminal. N-glycosylation sites scatter in the external surface, while no sites can be found in the embedded α-helixes bundle. The conserved N-glycosites hint that they may protect the important regions, or mutations in the RBD or cleavage sites can lead to zoonotic spillover and alteration of cell/tissue tropism, as exemplified by MERS and SARS CoVs. By analyzing their location in S-protein, the functionsof N-glycosylation sites mainly involve in: 1) Protein folding, 2) Protecting the antigen sites, 3) Protecting the S1/S2 cleavage site, 4) Protecting the C-terminal tail, 5) Affecting the receptor-ligand interaction^[60]^.

### 3.2 Glycan-binding participate in host invasion

Binding to host ligand is the earliest step during viral invading. The RBD (DPP4 for MERS CoV; ACE2 for SARS and SARS2 CoVs, 9-O-Ac-SA for OC43, HKU1 HCoVs, ANPEP (also known as CD13) for 229E CoV), or NTD (CEACAM1b for MHV CoV) of S protein on the surface of CoV bind to the receptor on the cell surface to facilitate the virus entering the host cell^[61]^. Therefore, whether an unknown glycan-binding mechanism participate in the SARS2 CoV interaction during invading remains unknown.

Previous studies from different CoVs have pointed out the glycan-binding is important for the enteropathogenicity^[62]^. What is more, the NTD of MHV, BCoV or OC43 CoV contain the same fold as human galectin domains, but with different binding ability. It also has described that the SA-binding domains located at the top of NTD, which consisted of β-sheets and two loops (Equal to the GCD1 in this study)^[32]^. Ruben’s work indicated this region was conserved in OC43-related family. The E182, W184, H185 and Y162 residues play the crucial roles in the SA-binding while the R143H, K181V, L186W, and I145T mutation didn’t alter the SA-binding ability^[63]^. In addition, a series of SA-related saccharides (including SA23Gal, SA26Gal, 3-sialyllactose or 6-sialyllactose) in the S-protein coordinations reveal MERS CoV adopt another strategy for SA binding (Similar to the GCD2 in this study). This SA binding domain located at the triangular tip of S-protein, which consisted of peripheral loops and small helixes. Obviously, the carboxyl group with S133, the hydroxyl groups at the C8,C9 atoms with K307, A92 as well as the Q36, I32 with the amide group form the multiple non-bonding interactions^[64]^.

Regardless of N-glyans, we speculated three potential GCDs in S-protein of SARS2 CoV by docking analysis, while two of three are similar to the above description. GCD1 lcateted at the top of NTD, which is consisted of one smaller helix, two samller β-sheets and long loops. Importantly, this domain showed either higher affinity to SA- or Gal-related glycans. Given the structural similarity between the NTD of SARS2, OC43, MERS or Galectin, it’s not hard to explain the affinity to Gal-related ligands of GCD1. Two loops and one helix of GCD2 formed a hydrophobic pocket, and V16, S254 and E154 are also conserved in Sarbecovirus subgenus. Similar location like the GCD3 in other CoVs has not been reported yet, we concluded this domain which locatedat the big groove of S-Protein side may be blocked by the nearby five N-glycans from the NTD, SD1, RBD and another nearby RBD. Although docking analysis provide powerful methods for ligands-receptor interaction, but the disadventages cannot be omitted, e.g. the docking is set to be occurred in the vaccum^[65]^.

In this study, the SA-related glycans has shown higher binding ability to these GCDs, it reflects the S-protein of SARS2 may have the potential ability for SA-binding. However, up to now, there are few validity reports from the anti-neuraminidase therapy. The SA-binding ability also could be proved by hemagglutination, such as the envelope glycoproteins from the paramyxoviridae, orthomyxoviridae or bunyaviridae[66-68]. Interestingly, there is another SA cleavaging enzyme, HE, in the confirmed SA-binding CoVs but missed in the SARS or SARS2 CoVs^[69–70]^. Though these indications may not support the SA-binding functions in the emerging SARS2 CoV. Considering the N-glycosylation is highly complicated, this protein-glycan interaction may be influenced by many factors.

More significantly, the GCD1 also showed higher affinity to Gal-related glycans, especially to the Galβ1-3GlcNAc structures. In our previous works of human saliva protein, the Galβ1-3GlcNAc-terminal structure was gradually accumulated along with chronic disease, such as the type 2 diabetes mellitus, hypertension or hepatocirrhosis^[71]^. It may help to explain that higher lethality in chronic patients due to dual binding mechanism.

### 3.3 N-glycosylation affects the host invasion in SARS2 CoV

CoVs use quite diverse strategies for interaction with cells involving the recognition of either specific protein receptors or certain derivatives of saccharides. Although the molecular mechanism of host-virus interaction in different CoVs have been described meticulously, however, the highly glycosylation in these interactions were commonly neglected.

Similar to S-protein, the HA in IVs and the gp160 in HIVs are the homotrimers and contain two subunits, e.g. the HA1 and HA2, gp120 and gp40 respectively^[72]^. It is well known that the SA-HA interaction play the crucial role in IVs, and the N-glycans at two N-glycosylation sites of HA even affect the host preference in H5N1 virus^[22,42]^. Considering the highly glycosylated in S-protein, it is not difficult to infer the viral invasion is affected by N-glycans directly or indirectly. As we point out, more than eight N-glycosylation sites distribute within a 50 Å radius of the center of ACE2-S-protein in SARS2 CoV (Figure S4), similar situations prevail in other CoV. Both 22 N-glycosylation sites could be found in S-protein monomer, 20 of 22 possess similar locations. The emerging N-glycosylation sites include the N149 in NTD and N657 near to S1/S2 cleavage site, while the miss N112 (N112S, equal to the N109 in SARS CoV) and N370 (T372A mutation in NSA result in the deletion of N-glycosylation site, equal to N357 in SARS CoV) are quite close to the RBD binding interface (Figure S5). The miss glycosylation sites near to the RBD result in a bigger exposed region and facilate the ACE2 binding during the viral invasion. Additionally, there are also more than ten N-glycosylation sites distribute around the binding interfaces from MHV-mCEACAM1a, MERS-DPP4 or others (Figure S6).

In previous analysis, we proposed several potential GCDs in the S-protein of SARS2 CoV, and the glycan-binding ability of the GCD to the N-glycan of ACE2 may facilitate the infection of host cells. During the SARS2 invasion, the RBD of S-protein bond to the N-terminal helix of ACE2, meanwhile, the N-glycan terminal (especially the SA and GalNAc residues) at N90 of ACE2 is apt to bind the GCD1 in S-protein, which may trigger the follwing conformational change of S-protein trimer. This hypothesis needs to be further verified.

## 4 Conclusion

The emerging SARS2 CoV results in tens of millions of infections and hundreds of thousands death in just a few months, the very contagious is higher than the known CoVs and attribute to the high affinity of its S-protein. This study has elaborated the distribution and functions of N-glycosylation sites in CoVs, as well as the potential GCDs in S-protein of SARS2 CoV. The high density of N-glycans surround the RBD-ACE2 interaction interface might suggest the dual binding mechanism, i.e. protein-protein interaction (RBD-ACE2) and glycan-protein interaction (N-glycan-GCD1) interactions. These results will help to explain the highly contagious of COVID-19.

## Acknowledgments

This work is supported by the National Natural Science Foundation (No. 31500130) and the emergency guidance fund for prevention of novel coronavirus pneumonia from northwest university (NWU002).

---

**Figure S1. Comparation of S-proteins from different CoVs.**

All the S-protein trimers are shown in cartoon mode and the N-glycosylation sites are denoted by yellow spheres. (A). SARS; (B). MERS; (C). MHV; (D). IBV; (E). PEDV; (F). NL63 CoV.

**Figure S2. Multi-Tyrosine residues in RBD participate in ACE2-RBD interaction.**

(A). Eight out of nine conserved Tyr residues constitute the hydrophilic pocket, similar binding mechanism also occurred in the SARS CoV. (B). The alignment of Tyr-rich region of RBD in Sarbecovirus subgenus and other Betacoronaviruses. All the contributing Tyr residues are labeled in red. SARS2 (NCBI NP:YP 009724390.1, Sarbecovirus), Bat SARS-like (AGZ48828.1, Sarbecovirus), SARS (ABD72993.1, Sarbecovirus), HKU9 (YP 001039971.1, Nobecovirus), HKU1 (ABD96188.1, Embecovirus), BCoV (ACB30202.1, Embecovirus), MERS (AID55093.1, Merbecovirus), OC43 CoV (AAX84792.1, Embecovirus).

**Figure S3. The superimposed NTDs and RBDs from different CoVs.**

Although the amino acids of S-protein varied greatly, the superimposed regions indicated they share the conserved core structures. (A) NTD; (B) RBD. Red: SARS; Blue: SARS2; Yellow: OC43; Cyan: MHV; Orange: MERS CoV.

**Figure S4. The N-glycosylation sites near to the RBD-ACE2 interaction interfaces.**

The N-glycosylation sites with the distances less than 50 Å are shown in red markers and yellow values. (A). One RBD in Up-standing state; (B). One RBD in lying state.

**Figure S5. The differences of N-glycosylation sites in the S-protein of SARS and SARS2 CoV.**

The N149 and N657 are only observed in SARS2 CoV while N112 and N370 are distinctive in SARS CoV.

**Figure S6. The protein-ligand binding complexs in the MERS-DPP4 and MHV-CEACAM1 interactions.** All the ligands are shown in red while the binding domains in S-proteins are shown in green. The N-glcosylation sites are shown in yellow spheres.

**File S1. The alignment of 1169 S-protein sequences from representative CoVs.**

**File S2. The procedure of force-filed file fabrication for glycoprotein MD simulation.**

**File S3. The N-glycosyltion sites of S-protein in Sarbecovirus members.**

**File S4. The N-glycosyltion sites of M-protein from representative CoVs.**

